# Langerhans cells and cDC1s play redundant roles in mRNA-LNP induced protective anti-influenza and anti-SARS-CoV-2 responses

**DOI:** 10.1101/2021.08.01.454662

**Authors:** Sonia Ndeupen, Aurélie Bouteau, Christopher Herbst, Zhen Qin, Zachary Hutchins, Drishya Kurup, Leila Zabihi Diba, Botond Z. Igyártó

## Abstract

Nucleoside modified mRNA combined with Acuitas Therapeutics’ lipid nanoparticles (LNP) have been shown to support robust humoral immune responses in many preclinical animal vaccine studies and later in humans with the SARS-CoV-2 vaccination. We recently showed that this platform is highly inflammatory due to the LNPs’ ionizable lipid component. The inflammatory property is key to support the development of potent humoral immune responses. However, the mechanism by which this platform drives T follicular helper cells (Tfh) and humoral immune responses remains unknown. Here we show that lack of Langerhans cells or cDC1s neither significantly affected the induction of PR8 HA and SARS-CoV-2 RBD-specific Tfh cells and humoral immune responses, nor susceptibility towards the lethal challenge of influenza and SARS-CoV-2. However, the combined deletion of these two DC subsets led to a significant decrease in the induction of PR8 HA and SARS-CoV-2 RBD-specific Tfh cell and humoral immune responses. Despite these observed defects, the still high antibody titers were sufficient to confer protection towards lethal viral challenges. We further found that IL-6, but not neutrophils, was required to generate Tfh cells and antibody responses.

In summary, here we bring evidence that the mRNA-LNP platform can support protective adaptive immune responses in the absence of specific DC subsets through an IL-6 dependent and neutrophil independent mechanism.

## INTRODUCTION

The vaccine platform based on the nucleoside-modified mRNA developed by Karikó and Weissman at the University of Pennsylvania (Karikó et al., 2005, 2011), combined with proprietary lipid nanoparticles (LNP) of Acuitas Therapeutics (Pardi et al., 2017), gained much attention with the ongoing SARS-CoV-2 pandemic. The platform was previously widely tested in animal models, and the studies reported induction of T follicular helper cells (Tfh) and robust protective antibody responses (Alameh et al., 2020; Pardi et al., 2018b). However, the immune mechanism by which this platform supports adaptive immune responses remains unknown. The nucleoside-modified and purified mRNAs do not induce strong inflammatory responses (Karikó et al., 2005, 2008, 2011). Still, the ionizable lipid component of these LNPs is highly inflammatory, causes rapid and robust neutrophil infiltration to the injection site, and was responsible for the development of the inflammatory responses characterized by the presence of high levels of inflammatory cytokines and chemokines (Ndeupen et al., 2021).

As professional antigen-presenting cells, dendritic cells (DCs) play critical roles in bridging innate and adaptive immune responses (Merad et al., 2013). DCs and DC-derived cytokines such as IL-6 are required to initiate the differentiation of naïve CD4^+^ T cells towards the Tfh cell lineage (Krishnaswamy et al., 2018). Using mice deficient in specific DC subsets, IL-6, or neutrophils, in combination with influenza and SARS-CoV-2 challenge models, here we show that the mRNA-LNP platform can support protective adaptive immune responses in the absence of specific DC subsets through an IL-6 dependent and neutrophil independent mechanism.

## RESULTS

### LCs and cDC1s show redundancy in driving anti-influenza and anti-SARS-CoV-2 responses triggered by the mRNA-LNP vaccine platform

The mRNA-LNP platform in which nucleoside-modified mRNA is combined with the proprietary LNPs of Acuitas Therapeutics drives effective adaptive immune responses in pre-clinical animal vaccine studies (Laczkó et al., 2020; Pardi et al., 2017). An LNP formulation with a different ionizable lipid from the same company is used in the Pfizer/BioNTech SARS-CoV-2 vaccine (Walsh et al., 2020). However, very little is known about the immune mechanism by which this platform supports the induction of Tfh cells and humoral immune responses. DCs, including Langerhans cells (LCs) and cDC1s, play essential roles in the induction of Tfh cells (Bouteau et al., 2019; Krishnaswamy et al., 2017; Lahoud et al., 2011; Levin et al., 2017; Yao et al., 2015). Therefore, here we tested the contribution of LCs and cDC1s in regulating adaptive immune responses triggered by this mRNA-LNP platform. Mice deficient in LCs (huLang-DTA, LC^-/-^; (Kaplan et al., 2005)), cDC1s (Batf3^-/-^, cDC1^-/-^; (Edelson et al., 2010)), or both (huLang-DTA by Batf3^-/-^, DKO; (Welty et al., 2013)), and littermate WT controls were intradermally immunized with 10 μg of mRNA-LNP coding for PR8 HA (Pardi et al., 2018a) as we previously described (Ndeupen et al., 2021). Seven and fourteen days later the Tfh and B cell responses were analyzed using flow cytometry in the skin draining lymph nodes. We found that in the absence of LCs or cDC1s, the Tfh and B cell responses specific to PR8 HA were comparable with WT levels (**Figure 1A-C**). However, when both DC subsets were missing, the Tfh and B cell responses were significantly reduced (**Figure 1A-C**). HAI analyses of the serum samples harvested 14 days post-immunization with PR8 HA, corroborated the flow cytometry data and revealed significant decrease in the DKO mice (**Figure 1D**). However, the total anti-HA serum IgGs determined by ELISA showed no major difference between WT and DC knockouts (**Figure 1E**).

**Figure 1.**
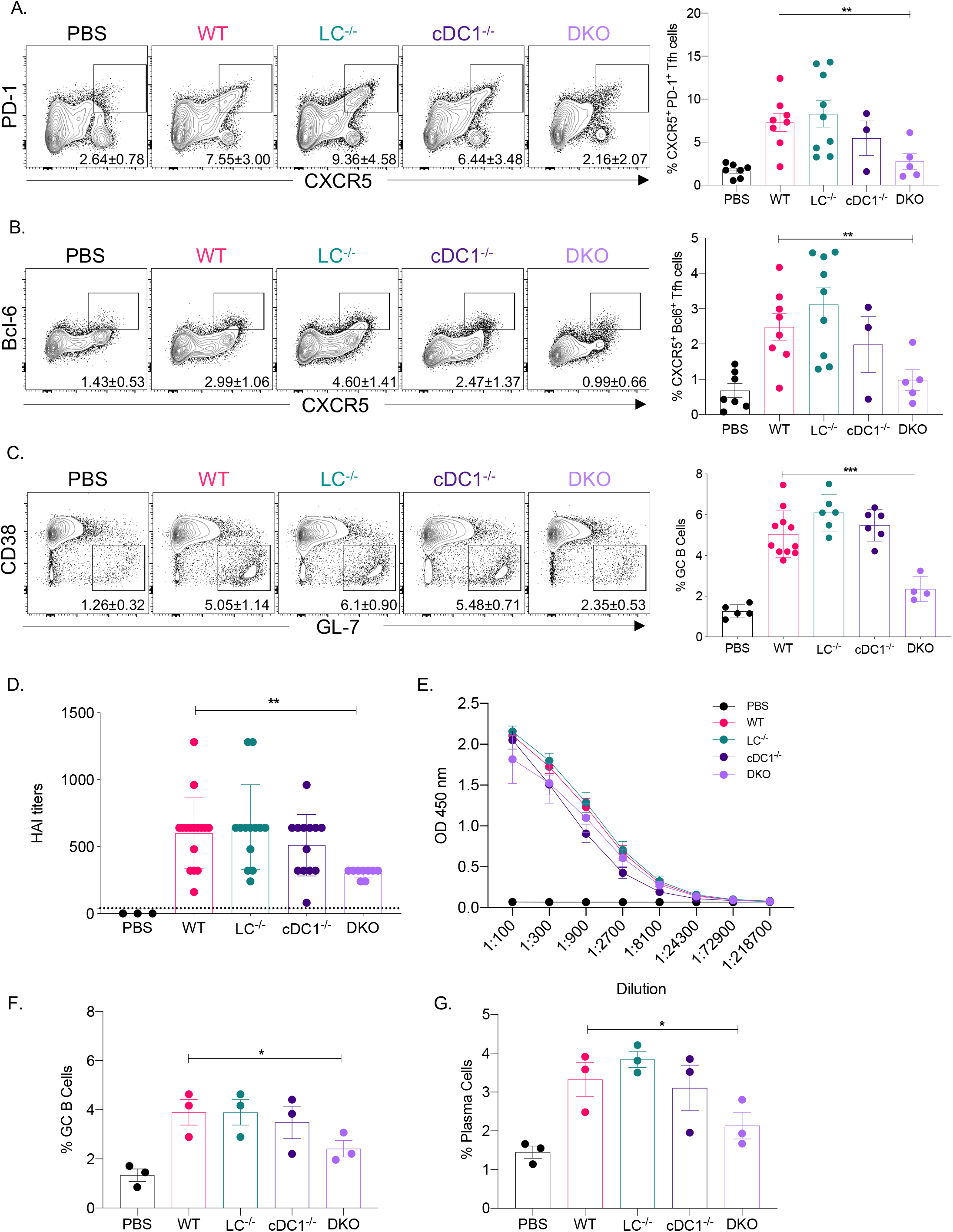
LCs and cDC1s play redundant roles in anti-flu and SARS-CoV-2 adaptive immune responses triggered by the mRNA-LNP platform. **A**. and **B**. The indicated mice were injected with 10 μg of mRNA-LNP coding for PR8 HA and the Tfh cell responses assessed in the skin draining lymph nodes 7 days post immunization. Mice injected with PBS served as control for basal levels. On the left representative flow plots while on the right summary graphs are presented. **C**. As in **A**, except that the lymph nodes were harvested 14 days post immunization and the GC responses determined using flow cytometry. **D**. and **E**. Serum samples from mice euthanized 14 days post immunization were assessed using HAI and ELISA. **F**. and **G**., as in **C**, except that the samples were from animals immunized with mRNA-LNP coding for SARS-CoV-2 RBD. For **A** through **E** data were pooled from 2-3 independent experiments. For **F** and **G**, one representative experiment. Each dot represents a separate animal. Two-tailed Student’s t test with Welch’s corrections were used to define significance. *p<0.05, **p<0.005, ***p<0.001.

To determine whether our findings can be generalized to other antigens, we immunized LC^-/-^, cDC1^-/-^, DKO, and littermate WT controls intradermally with 10 μg of mRNA-LNP coding for SARS-CoV-2 RBD (Laczkó et al., 2020). Fourteen days later harvested the skin draining lymph nodes and assessed the B cell responses using flow cytometry. The flow analyses showed that the RBD-specific B cell responses (GC and plasma cells), similarly to the influenza antigen, significantly decreased in the combined absence of LCs and cDC1s (**Figure 1F-G**). Thus, LCs and cDC1s play a redundant role in the induction of adaptive immune responses by the mRNA-LNP platform.

### Adaptive immune responses induced by the remaining APCs confer protection from lethal viral challenge

Mice lacking LCs, cDC1s, or both, still mounted WT levels of antigen-specific antibodies (**Figure 1E**). With PR8 HA immunizations, the HAI titers were magnitudes higher (**Figure 1D**) than the generally accepted protective levels (1:40). To test whether adaptive immune responses generated in the absence of specific DC subsets, confer protection against a lethal viral challenge, we repeated the experiments presented above. For the SARS-CoV-2 challenge studies, we bred the huLang-DTA (Kaplan et al., 2005) and Batf3^-/-^ (Edelson et al., 2010) mice to hACE-2 transgenic mice (Mccray et al., 2007). The generated genotypes were confirmed using PCR, and the hACE-2 positive and negative mice were further tested for virus susceptibility (data not shown). On day 14, post-immunization, we challenged the mice with lethal doses of PR8 influenza or SARS-CoV-2 intranasally. Unimmunized control mice (PBS), consisting of WT, and different DC knockouts dropped weight considerably. No significant differences were observed between the genotypes (**Figure 2A-B and Suppl. Figure 1A-B**), and were euthanized several days post-challenge (**Figure 2A-B**), while all the immunized groups, regardless of DC subset deficiency, remained protected (**Figure 2A-B**). The protective immunity was further confirmed for both influenza and SARS-CoV-2 by qRT-PCR on lung samples (**Figure 2C, and data not shown**). Thus, with the mRNA-LNP platform, in the absence of LCs and cDC1s, other APCs can drive protective adaptive immune responses.

**Figure 2.**
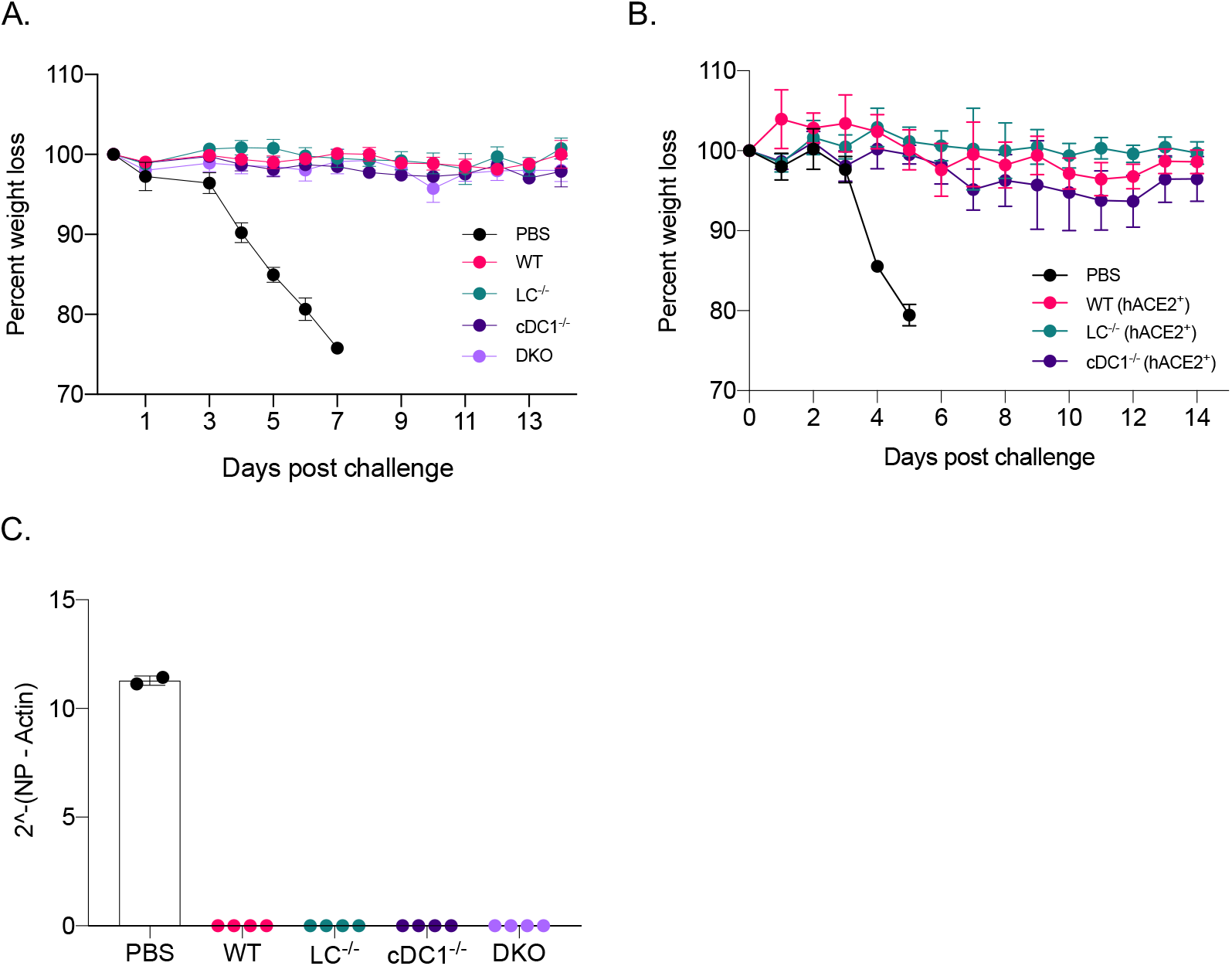
The mRNA-LNP platform supports the development of protective adaptive immune responses towards flu and SARS-CoV-2 even in the absence of certain DC subsets. **A.** The indicated animals were immunized with 10 μg of mRNA-LNP coding for PR8 HA or injected with PBS. Fourteen days later the animals were challenged with 5,000 TCDI PR8 influenza virus and the weight drop monitored as presented. Data from two independent experiments pooled, minimum 5 mice/group. **B**. As in **A**, but the indicated mice were immunized using 5 μg of mRNA-LNP coding for SARS-CoV-2 RBD and then 14 days later challenged with 10^5^ PFU SARS-CoV-2. One representative experiment with 5 mice/group is shown. **C**. Lung samples harvested 7 days post PR8 challenge were assessed using qRT-PCR for viral RNA. Each dot represents a separate animal.

### IL-6 is required for optimal induction of Tfh and B cell responses

IL-6 in mice in inflammatory settings is required for the induction of Tfh cells (Choi et al., 2013; Eddahri et al., 2009) and functions as a B cell growth factor (Kishimoto, 2010). We found that this mRNA-LNP platform induces high levels of IL-6 at the injection site (Ndeupen et al., 2021). Therefore, we tested whether IL-6 plays a role in adaptive immune responses triggered by the mRNA-LNP platform. Age-matched IL-6 knockout and WT mice were immunized intradermally with 10 μg of PR8 HA mRNA-LNP. The Tfh cells and the B cell responses were assessed by flow cytometry seven- and fourteen-days post-immunization. We found that in the absence of IL-6, both Tfh cell induction and B cell responses were significantly affected (**Figure 3A-C**). Immunized IL-6 deficient mice also showed decreased anti-HA serum IgG levels (**Figure 3D**). Thus, IL-6, similarly to other inflammatory models, plays a critical role in driving adaptive immune responses by the mRNA-LNP platform.

**Figure 3.**
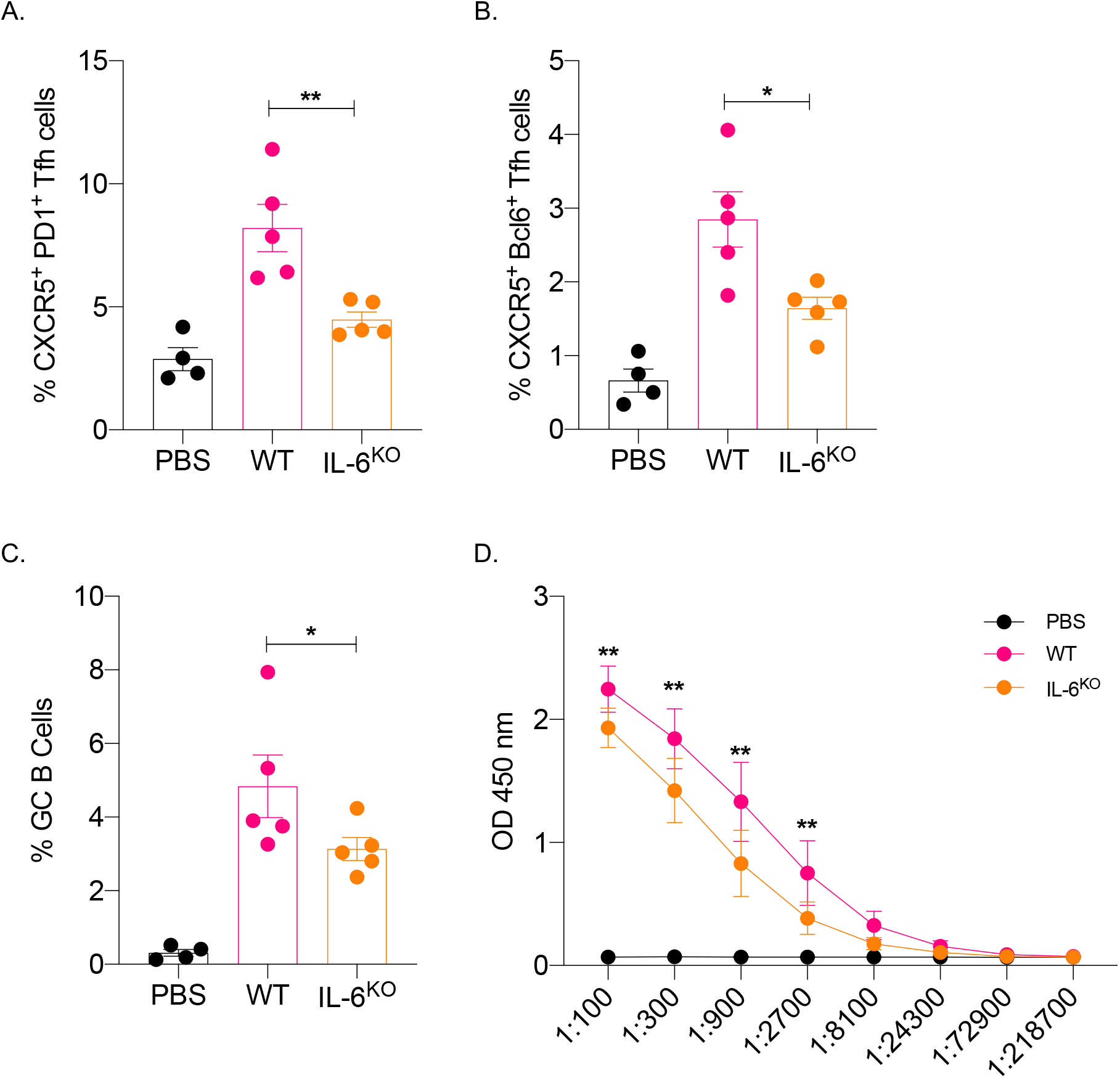
IL-6 is required for the induction of adaptive immune responses by the mRNA-LNP platform. **A**. and **B**. WT and IL-6 knockout mice were intradermally injected with 10 μg of mRNA-LNP coding for PR8 HA or PBS. Seven days later the Tfh cells were characterized in skin draining lymph nodes using flow cytometry. **C**. as in **A** and **B**, except the lymph nodes were harvested 14 days post injection and the GC B cell responses determined by flow cytometer. **D**. Serum samples at day 14 post injection were assessed for anti-HA IgG levels using ELISA. Data were pooled from two separate experiments. Each dot represents a separate animal. Two-tailed Student’s t test with Welch’s corrections were used to define significance. *p<0.05, **p<0.005.

### Neutrophils are dispensable for the Tfh and B cell responses induced by the mRNA-LNP platform

The mRNA-LNP platform through its ionizable lipid component triggers robust, rapid, but transient neutrophil infiltration at the delivery site (Ndeupen et al., 2021). Through induction of inflammatory milieu and activation of local APCs, and other cellular and molecular interactions, neutrophils can contribute to the generation of downstream adaptive immune responses (Van Gisbergen et al., 2005; Leliefeld et al., 2015; Li et al., 2019; Ludwig et al., 2006; Schuster et al., 2012). Therefore, to test the contribution of the neutrophils in regulation of adaptive immune responses triggered by the mRNA-LNP, we depleted them two days before immunization using 1A8 (anti-Ly-6G) antibody (Daley et al., 2008). Control mice were injected with isotype antibodies. The absence of neutrophils at the time and after immunization was confirmed using flow cytometry on blood cells (**Figure 4A-B and data not shown**). The mice were then immunized with 10 μg of PR8 HA mRNA-LNP as described above. Seven and fourteen days later, the Tfh cells and B cell responses, respectively, were assessed in the skin draining lymph nodes using flow cytometry. The absence of neutrophils did not significantly affect either PR8 HA-specific Tfh cell development or B cell responses (**Figure 4C-E**). The serum anti-HA IgG levels showed slight decrease in the absence of neutrophils, which however, did not reach significance (**Figure 4F**). Thus, these data suggest that neutrophils are unlikely to play a critical role in regulating adaptive immune responses induced by the mRNA-LNP platform.

**Figure 4.**
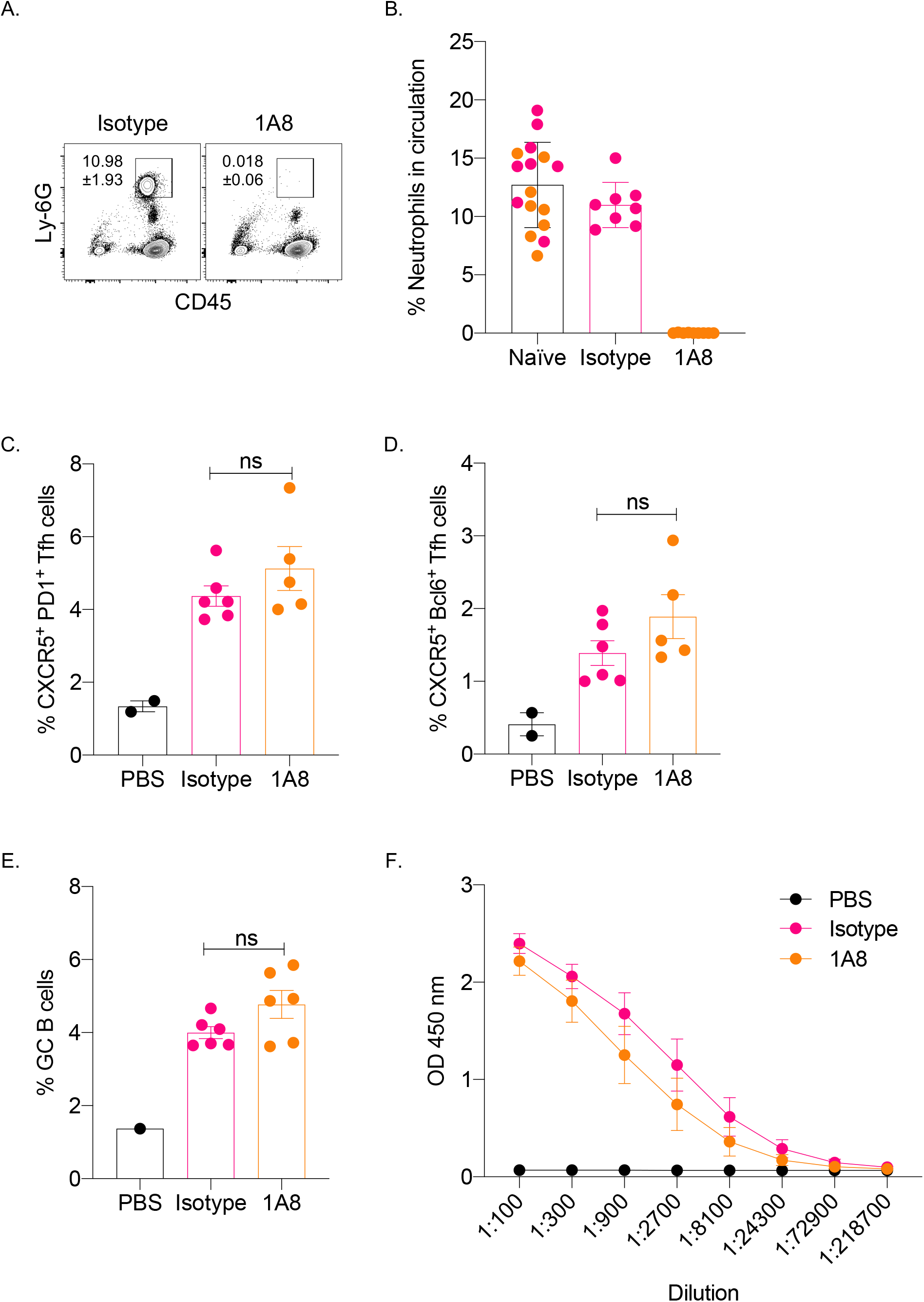
Lack of neutrophils at the time of immunization does not affect the induction of adaptive immune responses by the mRNA-LNP platform. **A**. and **B**. WT mice were either injected with isotype or neutrophil depleting antibodies (1A8) two days before immunization. **A**. Representative flow plot showing complete neutrophil depletion in the blood of 1A8 antibody injected mouse. **B**. Summary graph with multiple mice. Naïve indicates pre-injection levels of the neutrophils for the same mice. **C**. and **D**. Mice from **B** were intradermally injected with 10 μg of mRNA-LNP coding for PR8 HA or PBS. Seven days later the Tfh cells were characterized in skin draining lymph nodes using flow cytometry. **E**. as in **C** and **D**, except the lymph nodes were harvested 14 days post injection and the GC B cell responses determined by flow cytometer. **F**. Serum samples at day 14 post injection were assessed for anti-HA IgG levels using ELISA. Data were pooled from two separate experiments. Each dot represents a separate animal. Two-tailed Student’s t test with Welch’s corrections were used to define significance. ns = not significant.

## DISCUSSION

Here we show that the mRNA-LNP platform widely used in pre-clinical animal vaccine studies promotes protective adaptive immune responses against influenza and SARS-CoV-2 infections in the absence of LCs and cDC1s. We further identified IL-6 as a crucial inflammatory cytokine in supporting the induction of adaptive immune responses by this platform and showed that neutrophils were dispensable for these responses.

We recently described that the mRNA-LNP platform built on the nucleoside modified mRNA pioneered at the University of Pennsylvania by Karikó and Weissman (Karikó et al., 2005, 2011) combined with the proprietary LNPs of Acuitas Therapeutics is highly inflammatory (Ndeupen et al., 2021). The inflammatory property of this platform was likely due to its ionizable lipid component that led to activation of multiple inflammatory pathways and the production of a variety of different inflammatory cytokines and chemokines (Ndeupen et al., 2021). The inflammatory nature of the LNPs is crucial for the induction of adaptive immune responses because the mRNA alone did not induce detectable levels of adaptive immune responses (**Suppl. Figure 2**). However, it is unknown how the innate inflammatory responses induced by the LNPs lead to adaptive immune responses, especially to Tfh cell and B cell responses. DCs play a critical role in bridging the innate and the adaptive immune responses (Merad et al., 2013). We found that with the mRNA-LNP platform, LCs and cDC1s play redundant roles in driving Tfh cells and antibody responses. Nevertheless, the combined deletion of these DC subsets led to a significant decrease in the adaptive immune responses, confirming their essential roles in driving Tfh cells and B cell responses. However, even in the absence of these DC subsets, the mRNA-LNP platform induced antibody responses of WT levels and could confer protection from subsequent viral challenges. While the antibody titers showed no difference, the DKO-derived serum was less effective in an HAI assay. This suggests that the antibodies or some of the antibodies formed against HA in the absence of LCs and cDC1s might not effectively prevent agglutination but could still contribute to the protection, possibly by mediating ADCC (Gómez Román et al., 2014). Since the Tfh cell formation is severely affected in the DKO mice, it is possible that Th1 cells previously reported in the influenza settings (Miyauchi et al., 2016) could provide the necessary help for the B cell responses, which here might be of extrafollicular in nature. The antibody responses in this case, are likely driven by the cDC2s that are also potent in supporting antibody responses (Krishnaswamy et al., 2017). Since the platform is highly inflammatory, and the lipid-based nature of this platform allows the mRNA to reach and transfect cells far from the injection site, it is likely that even lymph node resident APCs could contribute to antigen presentation and regulation of adaptive immune responses. Antigen presentation by B cells is needed for the final maturation of Tfh cells (Crotty, 2019). B cells in the lymph node might also get transfected by the mRNA or have access to premade secreted antigens by other cells or presented to them by DCs. However, the details of the cellular and molecular interactions triggered by this platform remain to be elucidated.

Tfh cells and B cell responses in mice require the presence of IL-6 (Choi et al., 2013; Eddahri et al., 2009). DCs are thought to be the source of IL-6 (Cucak et al., 2009), and IL-6 is needed to initiate the Tfh cell program, including upregulation of the signature transcription factor Bcl-6 and CXCR5 (Chen et al., 2015; Choi et al., 2013). Our data support that IL-6 is also required for the induction of adaptive immune responses with this platform. However, further studies will be needed to identify the cellular source of IL-6. We observed high levels of IL-6 at the site of injection (Ndeupen et al., 2021). Still, cytokines present at the interface of the immunological synapse between APCs and naïve T cells are likely to be of higher importance (Benvenuti, 2016; Verboogen et al., 2016). The absence of IL-6 did not lead to a complete lack of Tfh cells and antibody responses. Therefore, other cytokines and co-stimulatory molecules previously described to play essential roles in regulating humoral immune responses, such as IL-1β and IFNα, are expected to contribute (Crotty, 2019; Krishnaswamy et al., 2018).

The immunological roles of DC subsets in SARS-CoV-2 infections are incompletely understood. Recent human studies have indicated that SARS-CoV-2 infections can affect DC biology (Benhnia, 2021; Campana et al., 2020; Saichi et al., 2021; Zhou et al., 2020). We found that naïve mice that lacked LCs or cDC1s and littermate WT controls succumbed to intranasal SARS-CoV-2 infections at the same rate (**Suppl. Figure 1B**). These data suggest that the absence of these DC subsets at the time of exposure does not render the host immunocompromised. More importantly, our vaccination studies showed a high degree of redundancy between DC subsets and cytokines. This vaccine platform can still induce protective antibody responses even in the absence of specific antigen-presenting cells and inflammatory cytokines. These are highly relevant findings and provide hope to individuals who might lack certain DC subsets or otherwise be immunocompromised to develop protective immune responses (Bigley et al., 2019).

From the perspective of that the mRNA-LNP platform is highly inflammatory and activates distinct inflammatory pathways and cells (Ndeupen et al., 2021), our findings that the mRNA-LNP vaccine can drive protective adaptive immune responses in the absence of certain innate immune cells and cytokines, are not surprising. While this might be a good thing from the vaccine’s perspective, we should tread carefully. Local, such as so-called Covid arm (Blumenthal et al., 2021), and distal inflammatory reactions, such as myocarditis, have been reported with the mRNA-LNP platform (Abu Mouch et al., 2021; Marshall et al., 2021; Montgomery et al., 2021; Shay et al., 2021). The robust inflammation driven by the ionizable lipid component of the LNPs (Ndeupen et al., 2021) likely contributed to the innate and adaptive immune reprogramming recently reported with this vaccine platform (Föhse et al., 2021). Thus, we think that more basic and translational research would be needed before this platform gets full approval for human use.

## ACKNOWLEDGMENTS

This work was supported by departmental start-up funds and by R01AI146420 to B.Z.I. We thank the flow cytometry core facility and the animal facilities for their help and assistance. Special thank you to Dr. Scott Hensley at UPenn for the PR8 (A/Puerto Rico/8/1934) influenza stock, and Dr. Holly Ramage at Thomas Jefferson University for the SARS-CoV-2.

## AUTHOR CONTRIBUTION

S.N. performed all the immunization experiments and analyzed the data. A.B. and C.H. performed the SARS-CoV-2 challenge experiments. Z.Q. determined the anti-HA antibody titers. Z.H. quantified the PR8 viral loads in the lung samples. D.K. and L.Z.D. did the SARS-CoV-2 dosing experiment on huACE-2 mice. B.Z.I. conceptualized the study, interpreted the data, and wrote the manuscript.

---

**Suppl. Figure 1. The absence of DC subsets at the time of exposure does not render the host immunocompromised. A.** The indicated naïve animals were challenged with 5,000 TCDI PR8 influenza virus and the weight drop monitored as presented. Data from two independent experiments pooled, minimum 5 mice/group. **B**. As in **A**, but naïve animals were exposed to 10^4^ PFU (sublethal dose) of SARS-CoV-2 and the weight drop monitored as presented. One representative experiment is shown with 5 mice/group.

**Suppl. Figure 2. mRNA-LNP dose titration experiment. A.** and **B.** WT animals were injected with increasing doses of mRNA-LNP coding for PR8 HA, or PBS or 10 μg of mRNA. Fourteen days later the B cell responses in the skin draining lymph nodes were determined using flow cytometry. **C**. Serum samples from **A** were assessed using HAI. Dotted line marks the 1:40 titer. Data from two independent experiments were pooled. Each dot represents a separate animal. Two-tailed Student’s t test with Welch’s corrections were used to define significance. *p<0.05, **p<0.005, ***p<0.001.

**Suppl. Figure 3. Gating strategy for identification of Tfh and GC B cells.**

**Suppl. Figure 4. SARS-CoV-2 dosing.** hACE2^+^ mice were inoculated intranasally with increasing doses of SARS-CoV-2 and weight drop recorded as presented. One representative experiment is shown with multiple mice.

**Suppl. Figure 5. Lower dose of mRNA-LNP still induces protective adaptive immune responses. A**. The indicated animals were immunized with 1 μg of mRNA-LNP coding for PR8 HA or injected with PBS. Fourteen days later the animals were challenged with 5,000 TCDI PR8 influenza virus and the weight drop monitored as presented. Data from two independent experiments pooled, minimum 5 mice/group.

## MATERIALS AND METHODS

### Mice

HuLang-DTA (Kaplan et al., 2005), Batf3^-/-^ (Edelson et al., 2010), huLang-DTA by Batf3^-/-^ (Welty et al., 2013) mice were previously published and bred in house. WT C57BL/6J, IL-6 KO and huACE-2 mice (Mccray et al., 2007) were purchased from Jax^®^ and bred in house. HuLang-DTA and Batf3^-/-^ mice were bred to huACE-2 mice in house. All experiments were performed with 6-15 weeks old mice. Mice were housed in microisolator cages and fed autoclaved food. Institutional Care and Use Committee approved all mouse protocols.

### Reagents and Viruses

For our studies, we used an LNP formulation proprietary to Acuitas Therapeutics described in US patent US10,221,127. These LNPs were previously carefully characterized and widely tested in preclinical vaccine studies in combination with nucleoside-modified mRNAs (Laczkó et al., 2020; Pardi et al., 2017, 2018a, 2018b). The following, previously published mRNA-LNP formulations were used: PR8 HA mRNA-LNP (Pardi et al., 2018a) and SARS-CoV-2 RBD mRNA-LNP (Laczkó et al., 2020). A/Puerto Rico/8/1934 influenza stock was a generous gift from Dr. Scott Hensley (University of Pennsylvania), while the SARS-CoV-2 was the original Washington isolate, and provided by Dr. Holly Ramage from Thomas Jefferson University.

### Intradermal immunization

Intradermal immunizations were performed as we previously described (Ndeupen et al., 2021). Briefly, the day before injections, the hair from the back skin of the mice was removed using an electric clipper, and then the site of injections wet-shaved using Personna razor blades. The next day the mice were injected intradermally with 2.5 μg/spot mRNA-LNP in 20 μl PBS (4 spots, 10 μg total) or PBS.

### Characterization of Tfh and B cell responses

At day 7 (peak of T cell responses; (Moon et al., 2009)) and 14 post-injections (peak of B cell responses; (Pape et al., 2011)), the mice were sacrificed and the skin draining lymph nodes (axillary, brachial and inguinal) harvested. Single-cell suspensions were generated using mechanical disruption through cell strainers. The mRNA-LNP platform does not require the use of T cell tetramers or fluorochrome-labeled antigen for B cells to study antigen-specific responses. There is only one antigen that the T and B cells react to, and the response is so robust that no need for magnetic enrichment either. Therefore, the cells were stained with either Tfh cell or B cell panels. The Tfh cell panel contained: fixable viability dye (Thermo Fisher), CD4 (GK1.5), CD44 (IM7), CD45 (OX-7), CD62L (H1.2F3), CD69 (H1.2F3), CXCR5 (L138D7), PD-1 (29F.1A12) and Bcl-6 (K112-91) from BioLegend and BD Biosciences. The gating strategy can be found in **Suppl. Figure 3A**. The B cell panel consist of dump (fixable viability dye, F4/80, CD11b), CD38 (90), B220 (RA3-6B2), CD138 (281-2), GL-7 (GL-7), Sca1 (D7), IgD (11-26c.2a) and IgM (RMM-1). The gating strategy is presented in **Suppl. Figure 3B**. The stained samples were run on LSRFortessa^™^ (BD Biosciences) and the resulting data analyzed with FlowJo 10.

### Intranasal challenge with PR8 influenza and SARS-CoV-2

The influenza dose used in these studies was previously described (Pardi et al., 2018a; Zens et al., 2016), while the SARS-CoV-2 dose was determined in house using hACE-2 mice (**Suppl. Figure 4**). Mice were anesthetized by intraperitoneal injection with a mixture of Xylazine/Ketamine and inoculated intranasally as follows. Mice received 5000 TCDI PR8 influenza virus or 10^5^ PFU SARS-CoV-2 (Washington isolate). Subsequently the mice were monitored daily for distress and weight loss. The weight loss data are presented as percent of original body weight.

### qRT-PCR for viral load

Mouse lung tissue was collected, and flash frozen prior to storage at −80°C. RNA was prepared using the RNeasy Micro kit (Qiagen, Cat: 74004), following the manufacturer’s instructions. 168 ng of RNA was reverse-transcribed using iScript Reverse Transcription Supermix for RT-PCR (Bio Rad, Cat: 1708840), following the manufacturer’s instructions. Quantitative PCR was performed with iTaq Universal SYBR Green Supermix (Bio Rad, Cat: 1725121), following the manufacturer’s instructions. Relative viral load was measured by ΔCT of PR8 influenza virus nucleoprotein (NP) using mouse β-Actin as a reference gene. Forward (5’) Pr8 NP: CAGCCTAATCAGACCAAATG, Reverse (3’): TACCTGCTTCTCAGTTCAAG. Forward (5’) β-Actin AGATTACTGCTCTGGCTCCTAGC and Reverse (3’): ACTCATCGTACTCCTGCTTGCT (Garcia et al., 2020; Li et al., 2021).

### ELISA

Nunc Immuno 96 well plates (Fisher Scientific) were coated with 1μg/ml (50 μl/well) HA protein (Sino Biological) diluted in carbonate/bicarbonate buffer (Fisher Scientific) overnight at 4 °C or 1 hour at 37 °C. After washing and blocking with TBS for 1 hour the serum samples were diluted and added to the plate. Serially diluted HA-specific monoclonal antibody (Sino Biological) served as standard. Anti-mouse IgG-HRP (1:20,000; Fisher Scientific) in combination with TMB (Fisher Scientific) solution was used for detection. The signals were read at 450 nm using accuSkan FC microplate photometer (Fisher Scientific).

### HAI

The assay was performed as previously described (Pardi et al., 2018a). Briefly, the sera were heat inactivated. Titrations were performed in 96-well, round-bottom plates (BD Biosciences). Sera were serially diluted twofold and added to four agglutinating doses of virus in a total volume of 100 μl. Turkey erythrocytes (Lampire Biological Laboratories) were added (12.5 μl of a 2% [vol/vol] solution). The erythrocytes were gently mixed with sera and virus, and agglutination was read after incubating for 1 h at room temperature. HAI titers were expressed as the inverse of the highest dilution that inhibited four agglutinating doses of influenza virus.

### Neutrophil depletion

Neutrophils were depleted using published protocol (Daley et al., 2008). Briefly, mice were either injected with 1 mg 1A8 (BioLegend) or isotype control antibodies (clone: RTK2758; BioLegend) intraperitoneally two days before immunization. Depletion efficiency was monitored in blood using flow cytometry.

### Statistical analyses

All data were analyzed with GraphPad Prism version 9.0.0. Statistical methods used to determine significance are listed under each figure.

