## Supplementary figures and images for "Langerhans cells and cDC1s play redundant roles in mRNA-LNP induced protective anti-influenza and anti-SARS-CoV-2 responses"

### Suppl. Figure 1.

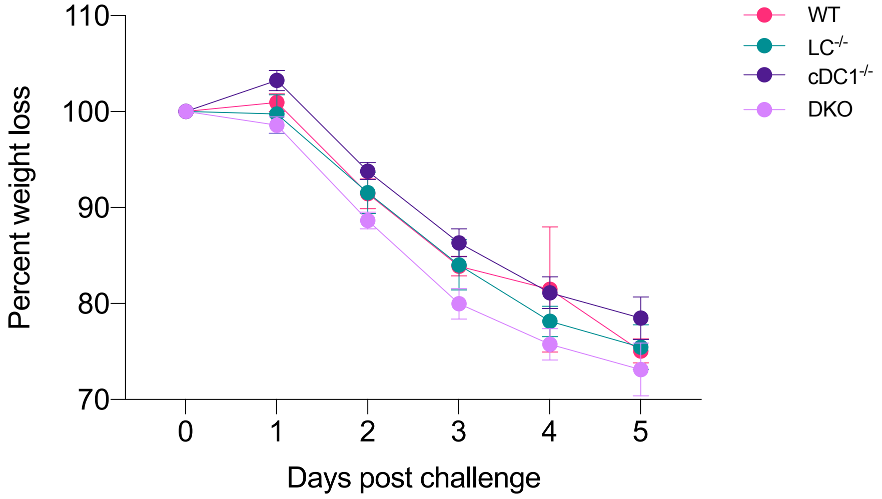

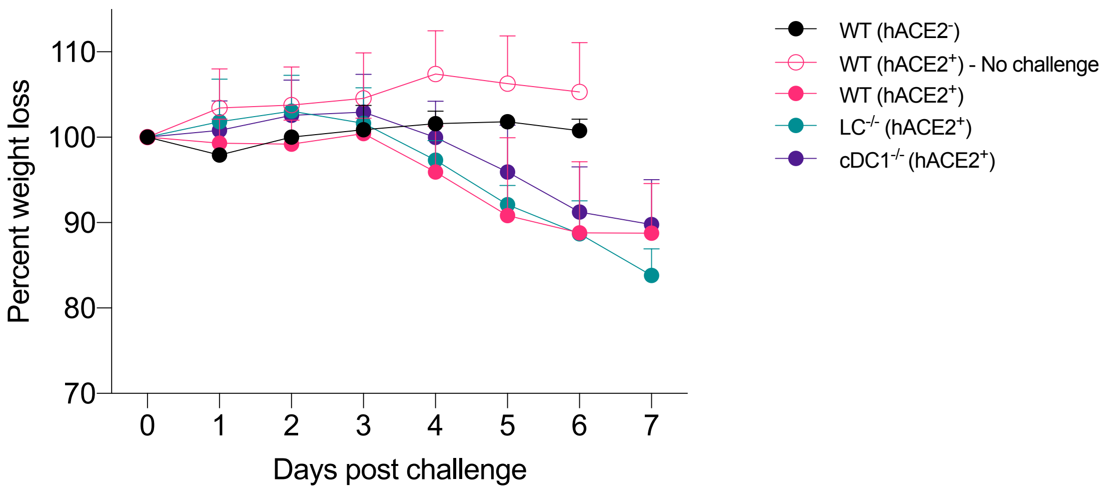

### Suppl. Figure 2.

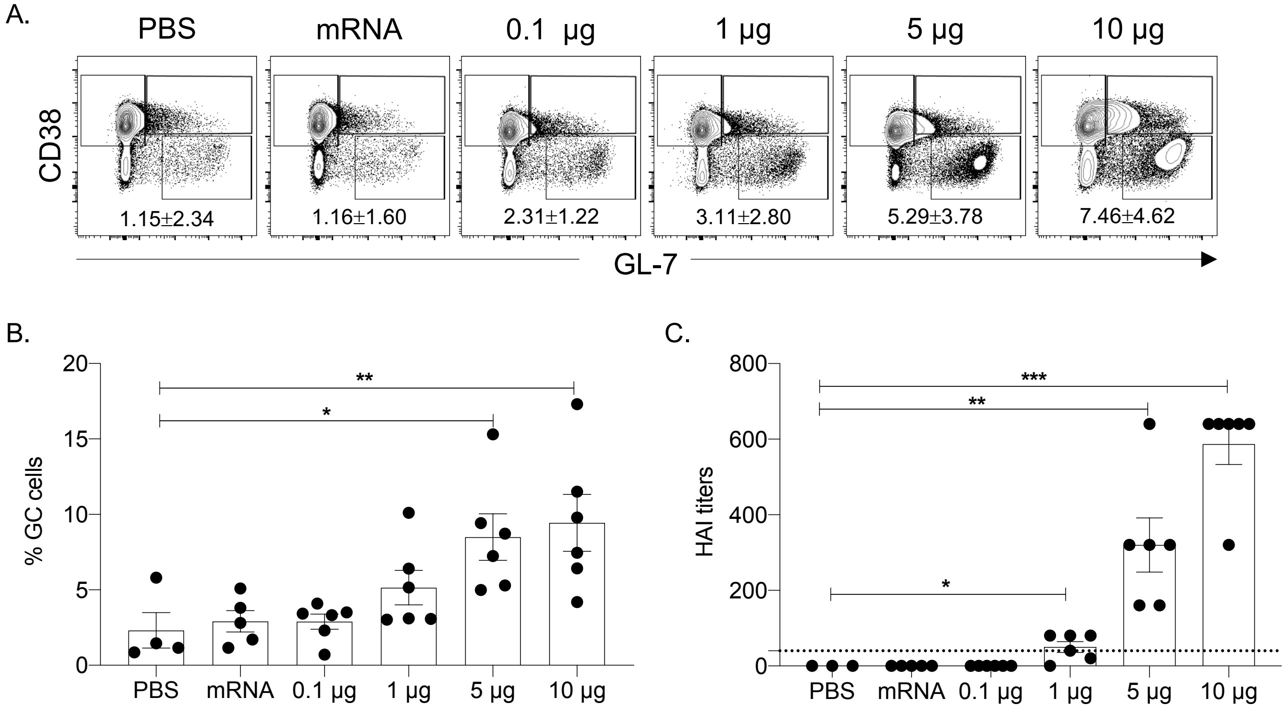

### Suppl. Figure 3.

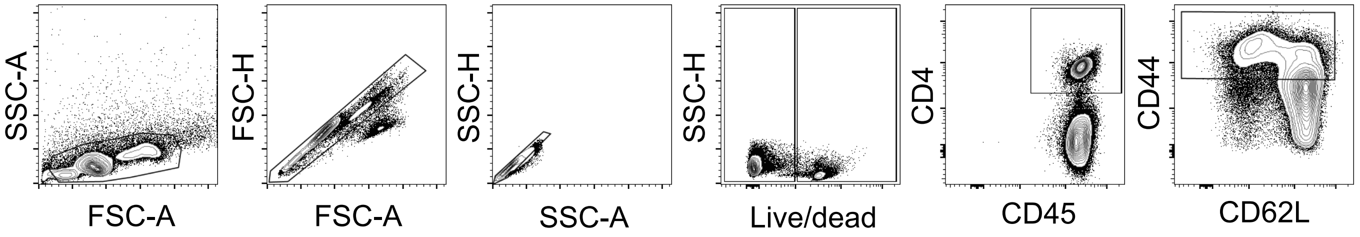

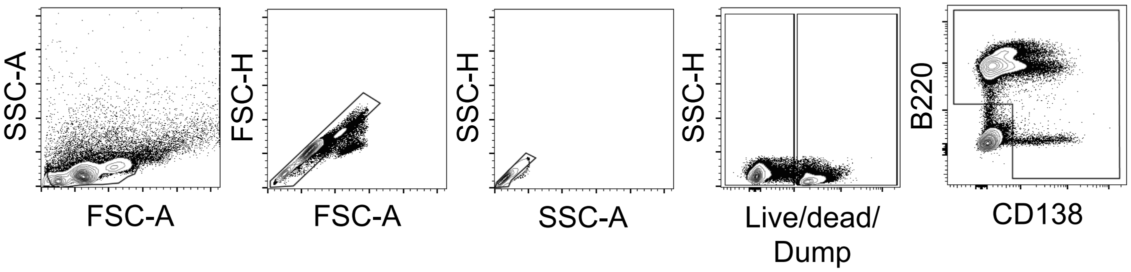

### Suppl. Figure 4.

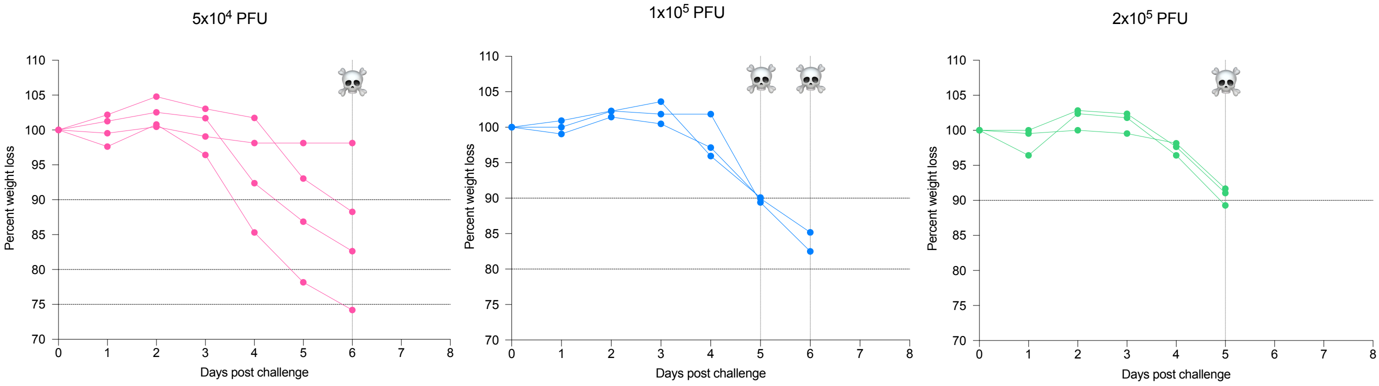

### Suppl. Figure 5

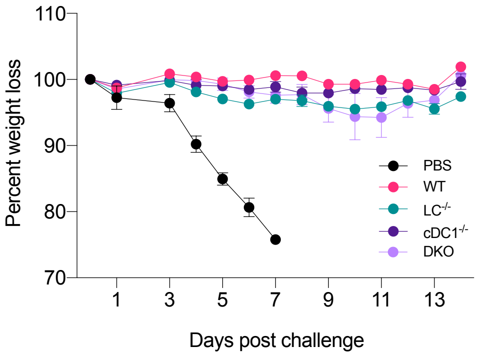
